# Effects of Centrally Acting Analgesics on Resting-State Electroencephalography Biomarker Candidates of Chronic Pain

**DOI:** 10.1101/2024.06.23.600313

**Authors:** Paul Theo Zebhauser, Felix Bott, Cristina Gil Ávila, Henrik Heitmann, Elisabeth S. May, Laura Tiemann, Enayatullah Baki, Thomas R. Tölle, Markus Ploner

**Author notes:** Corresponding author. Address: Department of Neurology, Technical University of Munich (TUM), Ismaninger Str. 22, 81675 Munich, Germany. (M. Ploner).

## Abstract

Resting-state electroencephalography (rsEEG) holds promise as a biomarker of chronic pain. However, the impact of centrally acting analgesics like opioids, antiepileptics, and antidepressants on rsEEG remains unclear. This confounds and limits the interpretability of previous studies and questions the validity of rsEEG biomarker candidates of chronic pain. We, therefore, aimed to elucidate the effects of opioids, antiepileptics, and antidepressants on common rsEEG biomarker candidates of chronic pain. To this end, we analyzed two large, independent rsEEG datasets, including 220 people with chronic pain. We performed preregistered multivariate Bayesian analyses and controlled for the potential confounds of age, pain intensity, and depression. The results predominantly provided evidence against effects of centrally acting analgesics on peak alpha frequency, oscillatory power in different frequency bands, and connectivity-based network measures. Although these findings do not rule out any effects of analgesics on rsEEG, they argue against medium to large effects of centrally acting analgesics on rsEEG. Thus, the findings strengthen the validity of rsEEG biomarker candidates of chronic pain and might thereby help to develop clinically valuable biomarkers of chronic pain.

## 1. Introduction

Establishing biomarkers is a crucial challenge to improve the diagnosis and treatment of chronic pain (CP) [4]. As brain function is altered in CP [16], measures of brain function could help to develop such biomarkers [18]. Electroencephalography during the resting state (rsEEG) is a broadly available and affordable measure of brain function that is scalable to large populations [20]. Hence, many studies have tried identifying EEG biomarkers for predicting, diagnosing, and monitoring CP and treatment responses. A recent systematic review showed higher power of brain activity at theta (4-8 Hz) and beta (13-30 Hz) frequencies in people with CP compared to healthy participants [35]. Moreover, evidence for slowing of the peak frequency of alpha oscillations (8-13 Hz) has been found [35]. Beyond, connectivity and network measures are promising rsEEG biomarker candidates for CP [17; 25].

Validity and specificity are significant challenges in biomarker development [4]. Many people with CP take centrally acting analgesics, which might affect EEG measures of brain function and thereby limit the validity and specificity of EEG biomarkers of chronic pain. Thus, understanding the effects of centrally acting analgesics is crucial for developing EEG biomarkers. Centrally acting analgesics mainly comprise opioids (OPDs), analgesic antiepileptics (AEDs), and analgesic antidepressants (ADs) [3]. Evidence for their effects on EEG is scarce and inconsistent, as most studies have been conducted in small samples of healthy participants. Furthermore, in studies of people with CP, medication is often not reported [35]. Concerning OPD, a few studies in healthy participants found power increases at delta (0.1-4 Hz) and theta frequencies [1; 19]. In people with CP, alpha and beta power increases were seen after OPD administration [9]. Concerning the AEDs gabapentin and pregabalin, a study in healthy participants observed slowed peak alpha frequency [26], and a study in people with CP [11] found increased theta power. Regarding ADs, serotonin and norepinephrine reuptake inhibitors (SNRI) and tricyclic antidepressants (TCA) are most frequently used in pain medicine [3; 8]. In healthy participants, one study found increases in theta and beta power [19]. To our knowledge, no studies have been conducted on AD effects on rsEEG in people with CP. Thus, whether and how centrally acting analgesics affect rsEEG measures in people with CP is not fully clear.

Here, we aimed to determine the effects of the most frequently used centrally acting analgesics on rsEEG biomarker candidates of CP. To test the replicability and generalizability of the findings, we analyzed two independent large EEG datasets recorded with different EEG devices. We analyzed measures of oscillatory power at various frequencies and peak alpha frequency, as these measures are the rsEEG biomarker candidates that have been most frequently analyzed in CP [35]. In addition, we analyzed the effects of connectivity-based network measures of brain activity, which have been put forward as biomarker candidates in CP and neuropsychiatry [17; 25]. The overarching aim of this approach is to further the development and interpretation of EEG-based biomarkers of CP.

## 2. Methods

### 2.1. Sample characteristics

To increase the validity and generalizability of findings, we analyzed EEG data of two independent samples of people with CP recorded with different EEG devices. Sample A comprised 118 people with CP from previous studies [12; 31] and EEG was recorded using 64 wet electrodes. Sample B comprised 102 people with CP from an ongoing study on EEG correlates of neuropsychiatric symptoms in CP (*clinical*.*trials*.*gov-identifier NCT05261243*) and EEG was recorded using 29 electrodes. Three EEG recordings from sample B had to be excluded from analyses for data quality reasons (excessive artifacts), resulting in a final sample size of *n=99*. Table 1 shows the demographics and clinical characteristics of the participants of both studies, which were comparable between samples. For both studies, we included people with the diagnosis of a CP condition treated at our interdisciplinary outpatient clinic for pain (convenience sampling); the frequency and type of diagnoses are shown in Table 1. Exclusion criteria were identical for studies (severe concomitant neurological or psychiatric condition, primary headache condition, age under 18 years, inability to sign informed consent for study participation due to language barrier or other reasons). To limit potential confounds of other substance classes on EEG activity, we excluded people with CP taking benzodiazepines (*n=2*) or Z-substances (*n=3*) [33], cannabinoids (*n=9*) [29; 30], neuroleptics (*n=1*) [13], and AEDs other than gabapentin or pregabalin (*n=3*) [21] from participation in the study. All participants provided written informed consent, and study protocols were approved by the Ethics Committee of the Medical Faculty of the Technical University of Munich (TUM) and conducted following relevant guidelines and regulations.

**Table 1.** Clinical and demographic variables of the participants. For continuous variables, mean ± standard deviation are reported, f=female, m=male, CP=chronic pain, CBP=chronic back pain, NPP=neuropathic pain, WSP=widespread pain, JP=joint pain, NRS=numeric rating scale, BDI=Beck’s Depression Inventory, *note that different ranges apply for the BDI *(0-63 used in Sample A)* and the PROMIS-29-subscale for depressive symptoms *(4-20, used in Sample B)*. For three cases from sample B, information on depression and pain scores was lost due to technical problems. As these variables affected only covariates, mean imputation was performed.

|  | Sample A | Sample B |
| --- | --- | --- |
| n (included in analysis) | 118 | 99 |
| Age (years) | 56.6 ± 15.9 | 57.0 ± 17.4 |
| f:m | 76:42 | 59:40 |
| Primary CP diagnosis | CBP n=74<br>NPP n=20<br>WSP n=11<br>JP n=13<br>Other=0 | CBP n= 39<br>NPP n= 27<br>WSP n= 3<br>JP n= 18<br>Other= 12 |
| Pain intensity (NRS 0-10) | 5.5 ± 1.8 | 5.5 ± 2.2 |
| Pain duration (months) | 111.3 ± 104.9 | 83.4 ± 101.9 |
| Depressive Symptoms (BDI-II/PROMIS-29)* | 13.9 ± 8.7<br>(BDI-II) | 8.6 ± 3.2<br>(PROMIS-29) |

### 2.2. Medication

All people with CP included in this study were treated at the Center for Interdisciplinary Pain Medicine of the TUM. Detailed medication plans, including specific substances and doses, were acquired from patient files and cross-validated on the day of the EEG recording. To investigate dose-dependent effects on EEG variables, we calculated oral morphine equivalents (milligrams/day) based on recent guidelines [7; 34]. Similarly, we calculated pregabalin equivalents for gabapentin (milligrams/day) with a 1:6 ratio [2]. In the case of PRN medication *(“pro re nata”*), daily averages were calculated and added to basic medication doses. Regarding ADs, we combined substance classes (with few exceptions, the SNRI duloxetine or the TCA amitriptyline were used) regardless of their mode of action. Because of the overall low number of people with CP taking ADs, subgroup analyses were not feasible.

### 2.3. EEG recordings

For EEG recordings of both samples, participants were seated in a comfortable chair in a quiet room and were instructed to stay in a wakeful and relaxed state. 5-minute segments of eyes-closed and eyes-open were recorded. Eyes-closed data was used for all analyses.

For sample A, data was recorded using a wet EEG system with 64 electrodes (EasyCap, Herrsching, Germany) and a BrainAmp MR plus amplifier (Brain Products, Munich, Germany). The electrode layout included all 10-20 system electrodes and additional electrodes Fpz, CPz, POz, Oz, Iz, AF3/4, F5/6, FC1/2/3/4/5/6, FT7/8/9/10, C1/2/5/6, CP1/2/3/4/5/6, TP7/8/9/10, P1/2/5/6/7/8, and PO3/4/7/9/10. During recordings, electrodes were referenced to FCz and grounded at AFz. Data were sampled at 1000 Hz (0.1 mV resolution) and impedances were kept below 20 kΩ.

For sample B, data was recorded using a dry EEG system with 29 electrodes (CGX Quick 32-r, San Diego, CA) with a built-in wireless amplifier. The electrode layout included all 10-20 system electrodes and additional electrodes Fpz, AF7/8, FC5/6, CP5/6, P07/8, and Oz. During recordings, electrodes were referenced to and grounded at A1 (left earlobe). Data were sampled at 500 Hz (0.2 mV resolution), and impedances were kept below 2500 kΩ.

### 2.4. EEG analysis

The complete analysis plan was preregistered on osf.io: https://osf.io/uzgxw. All EEG data were preprocessed and analyzed in MATLAB using the openly available pipeline *DISCOVER-EEG* [10] with the EEGLab [6] and Fieldtrip [22] toolboxes. Preprocessing included line noise removal, bad channel rejection, re-referencing, independent component analysis, bad component removal, bad channel interpolation, and bad time segment removal. For all analyses, frequency bands were defined according to the *COBIDAS MEEG* recommendations [23] as follows: theta (4 to < 8 Hz), alpha (8 to < 13 Hz), beta (13 to 30 Hz), and gamma (> 30 to 80 Hz). We extracted the following EEG features: peak alpha frequency (measured as the center of gravity, or the local maximum of the alpha band), power spectra (global power), and three global connectivity-based network measures (see below). Power spectra were computed for every channel and averaged across epochs and channels to obtain a single global power spectrum. Absolute power was calculated by summing up power values for frequency bands, and averages for frequency bands were determined for statistical analyses. Relative power was calculated by normalizing absolute power for respective frequency bands by total power values across frequency bands. To evaluate connectivity-based network measures, we incorporated one phase-based and one amplitude-based measure of functional connectivity, as these approaches capture at least partly different neuronal mechanisms [28]. We selected the debiased weighted Phase Lag Index (dwPLI) and the orthogonalized Amplitude Envelope Correlation (AEC), which are commonly used in EEG analysis. These measures are undirected and have low susceptibility to spurious volume conduction effects. We performed source reconstruction of the preprocessed data. We applied an array-gain Linear Constrained Minimum Variance (LCMV) beamformer to project the data to source space. We used centroid regions of the Schaefer atlas with 100 parcels [27] as a source model. The lead field was modeled using a volume conduction model based on the Montreal Neurological Institute (MNI) template available in FieldTrip (standard_bem.mat) and the source model. Spatial filters were constructed using covariance matrices from the band-pass filtered data and the specified lead fields. To address rank deficiencies in the covariance matrix, a 5% regularization parameter was applied. DwPLI and AEC were computed for each frequency band and combination of 100 reconstructed virtual time series. We derived three global graph theory-based network parameters (global clustering coefficient, global efficiency, and smallworldness) for each frequency band and method of connectivity calculation (dwPLI, AEC). The global clustering coefficient is a network measure of functional segregation and is defined as the average clustering coefficient of all nodes. Global efficiency is a measure of functional integration in a network and is defined as the average of the inverse shortest path length between all pairs of nodes. Smallworldness compares the ratio between functional integration and segregation in the network against a random network of the same size and degree. All network parameters were computed on thresholded and binarized connectivity matrices (binarized by keeping the 20% strongest connections).

### 2.5. Statistical analyses

For all statistical analyses, Bayesian analyses were performed using JASP Version 0.18.3.0 [14]. Bayesian analyses were particularly suitable to answer our research questions, as they also enable quantification of evidence for the absence of an effect (‘negative results,’ i.e., no effects of centrally acting analgesics on rsEEG biomarkers candidates). The Bayes factor (BF_10_) was used to quantify effects *for* (possible “evidence for absence”) or *against* the null hypothesis. BF_10_ was interpreted as strong, moderate, or weak evidence (>10, 3-10, >1<3 regarding the alternative hypothesis; <; 0.1, >0.1<0.33, 0.33-1 regarding the null hypothesis) according to recent guidelines [32]. All analyses were conducted with default priors due to the absence of prior statistical evidence regarding the effects of respective substance classes on EEG variables.

We pursued a stepwise approach to analyze the effects of centrally acting analgesics on rsEEG features. For all models, age, pain intensity, and depressive symptoms were entered as covariates, as associated confounding effects on rsEEG features are plausible. Non-normally distributed variables were BoxCox-transformed for all statistical models.

First, we performed group comparisons. To analyze differences in EEG features between people with CP taking centrally acting analgesics and those who did not, we used Bayesian ANCOVAs with a binary coded independent variable (IV) encoding the group and continuous dependent variable (DV) representing the EEG feature of interest (e.g., peak alpha frequency). To assess the potential effects of different substance classes, we used Bayesian ANCOVAs with 5-level IVs (no centrally acting analgesic, OPDs only, AEDs only, ADs only, or any combination of these substance classes). For all ANCOVAs, the main effects of the IV were analyzed (BF_10_ for inclusion in the model).

Second, we investigated dose-dependent effects. To evaluate the dose-dependent effects of OPDs and AEDs, we applied Bayesian regression models with either oral morphine equivalents (milligrams/day) or pregabalin equivalents (milligrams/day) as continuous predictors and EEG features as DVs. For all regression models, the effects of OPDs or AEDs were analyzed (BF_10_ for inclusion in the model).

Lastly, we examined cumulative analgesic use. This was assessed with the Medication Quantification Scale 4.0 (MQS) [5] for all people with CP using an analgesic (including peripheral analgesics, *n=88/70* for samples A and B, respectively). The MQS is an established tool to quantify the overall risk associated with the use of various analgesics, including central nervous system effects [5]. We analyzed effects on EEG features using Bayesian regression models (MQS score and covariates age, pain intensity, depression score as predictors, rsEEG biomarkers as DV). Equivalent to regression models for dose-dependent effects, BF_10_ for inclusion in the model was interpreted for MQS scores.

### 2.6. Data and code availability

Data and code are available online without restrictions. Datasets are available at https://osf.io/uzgxw. Data is stored in EEG-BIDS format [24]. Code for analysis of EEG data is accessible through *DISCOVER-EEG* [10], an openly available preprocessing and analysis pipeline for EEG biomarker discovery in clinical neuroscience.

## 3. Results

### 3.1. Use of centrally acting analgesics

In sample A, 75 of 118 (63.6%) people with CP took at least one centrally acting analgesic. Forty-three people with CP (36.4 %) took an AED, 35 (29.6 %) took an OPD, and 33 (28.0 %) took an AD as part of their daily pain medication. In sample B, 61 of 102 (59.8%) people with CP took at least one centrally acting analgesic, of whom 35 people with CP (34.3 %) took an AED, 30 (29.4 %) took an OPD, and 31 (30.4 %) took an AD. Fig. 2 illustrates the number of people with CP taking the different substance classes and their overlap in the case of multiple substance classes for samples A and B.

**Fig. 1.**
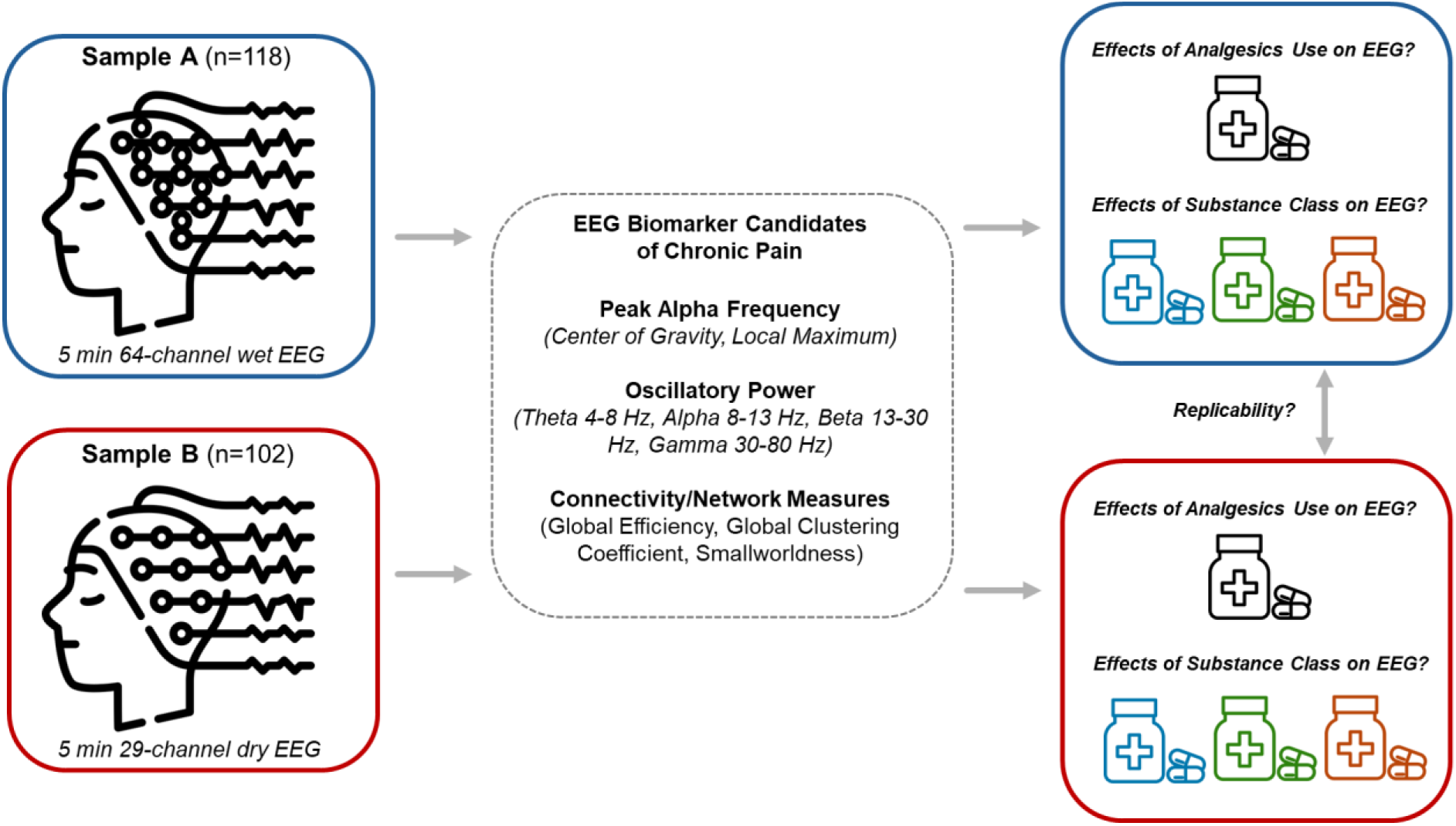
shows an overview of the study design and the research questions.

**Fig. 2:**
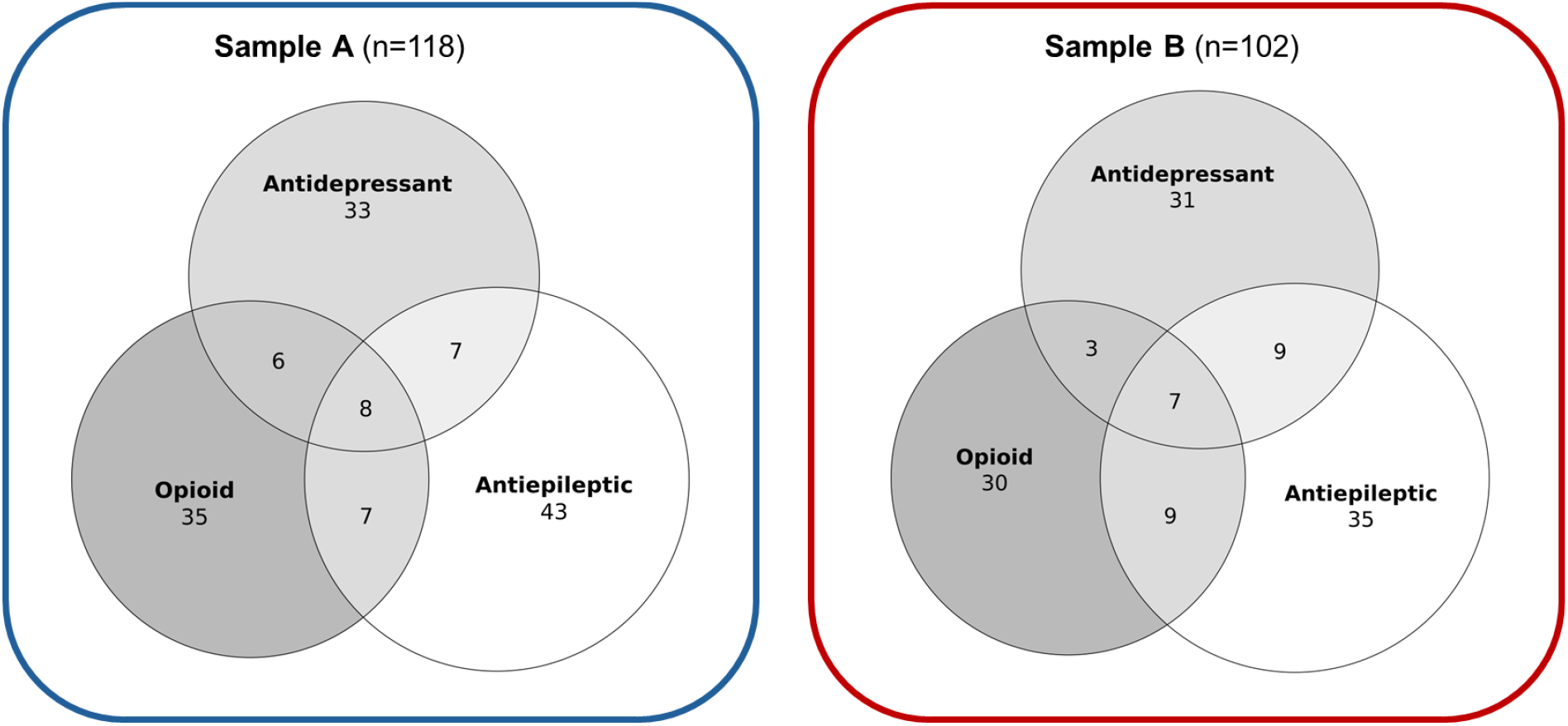
Venn diagrams showing patient numbers with daily intake of centrally acting analgesics. In sample A, n=43, and in sample B, n=41 people with CP did not use any centrally acting analgesic (not shown in figure).

### 3.2. Effects of centrally acting analgesics on EEG features

#### 3.2.1. Group comparisons

We first analyzed differences in rsEEG biomarker candidates between participants taking any centrally acting analgesics and those who did not. To this end, we performed Bayesian ANCOVAs (2-level IV; any vs. no centrally acting analgesic; Fig. 3, upper left and right panels) for both samples. In sample A, we found mostly moderate evidence against differences between groups (Fig. 3, upper left panel). Very few exceptions were found regarding network measures (mainly in the gamma band), for which we observed very weak evidence for group differences (*BF*_*10*_ = 1.06 < 1.50). None of these effects were found in sample B. In sample B, we also observed mostly evidence against group differences (Fig. 3, upper right panel). We did, however, find moderate evidence for a higher global clustering coefficient (*BF*_*10*_ *= 3*.*03*) and higher smallworldness (*BF*_*10*_ *= 3*.*58*, each based on AEC) in the theta band and higher relative alpha power (*BF*_*10*_ *= 4*.*78*) in participants taking centrally acting analgesics. These effects were not found in sample A.

**Fig. 3:**
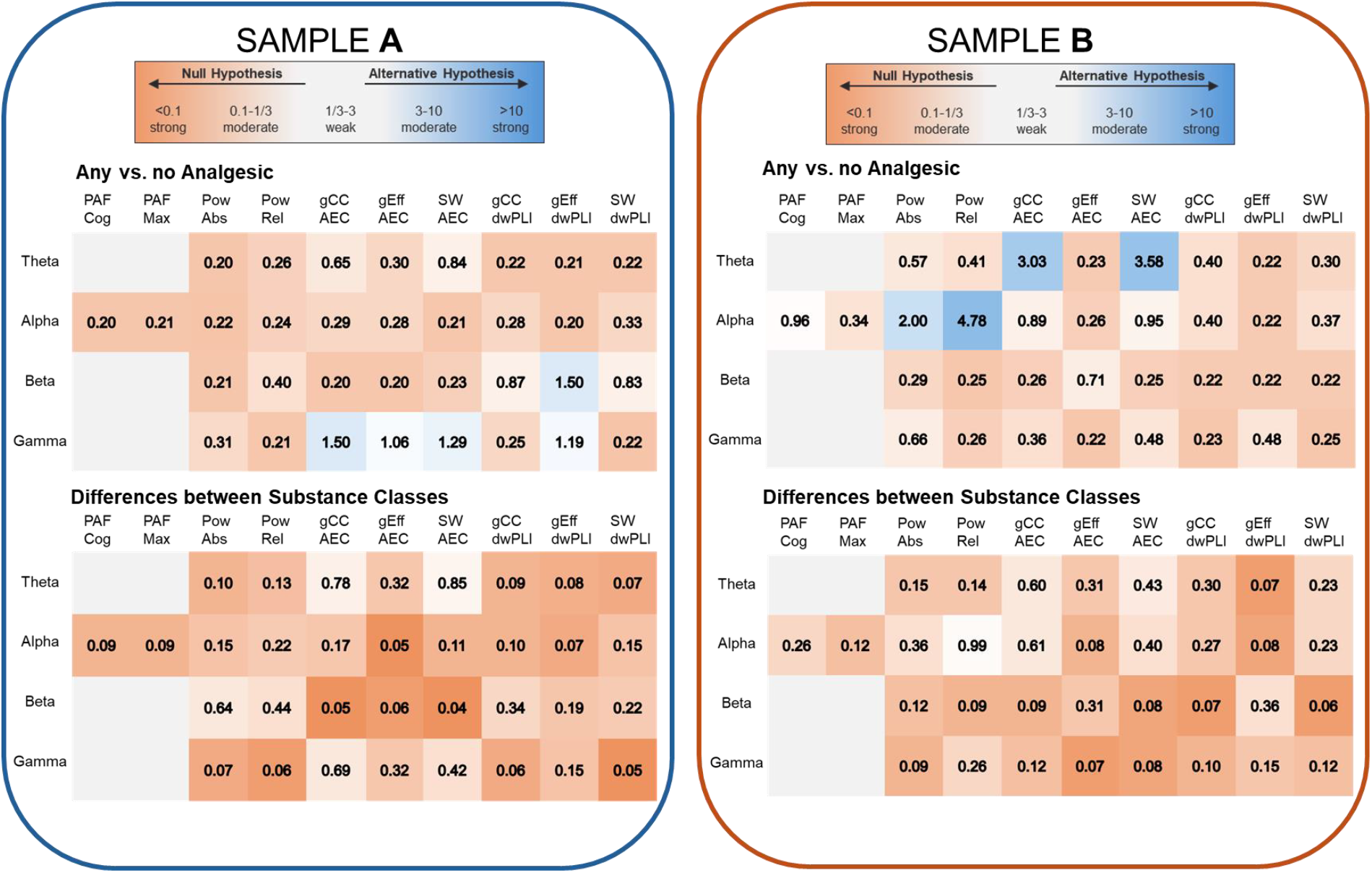
Group comparisons of EEG features in people with CP split by centrally acting analgesics use. Overview of BF10 resulting from ANCOVAs comparing rsEEG features between people with CP using centrally acting analgesics and people with CP not using centrally acting analgesics (upper matrices) and between people with CP taking medications from different substance classes (opioids, antiepileptics, antidepressants; combination thereof, bottom matrices). For all models, age, pain intensity, and depressive symptoms were included as covariates. BF=Bayes Factor, PAFCog=peak alpha frequency (Center of Gravity), PAFMax=peak alpha frequency (Local Maximum), PowAbs=Absolute Power, PowRel=Relative Power, gCC=global Clustering Coefficent, gEff=global Efficiency, SW=Smallworldness, AEC=Amplitude Envelope Correlation, dwPLI=debiased weighted Phase Lag Index.

We next assessed differences in rsEEG biomarker candidates between participants taking different substance classes. To this end, we performed Bayesian ANCOVAs with 5-level IVs (no centrally acting analgesic, OPDs, AEDs, ADs, or any combination thereof; Fig. 3, lower left and right panel). The results showed mostly moderate evidence against group effects in both samples. None of the comparisons yielded evidence in favor of a group difference.

Thus, the pattern of group comparisons did not provide consistent evidence in favor of, but mostly evidence against effects of centrally acting analgesics on rsEEG biomarker candidates.

#### 3.2.2. Dose-dependent effects

We further assessed dose-dependent effects of OPDs and AEDs. To this end, we applied regression models with either oral morphine equivalents (milligrams/day) or pregabalin equivalents (milligrams/day) as continuous predictors and rsEEG features as DVs in both samples. Almost exclusively, these analyses revealed evidence against OPD and AED doses as predictors of rsEEG features (see Fig. 4). The only exception was moderate evidence (*BF*_*10*_ *= 4*.*73*) for a dose effect of OPDs on global efficiency in the theta band (the higher the dose, the lower the global efficiency) in sample A. However, this finding was not replicated in sample B.

**Fig. 4:**
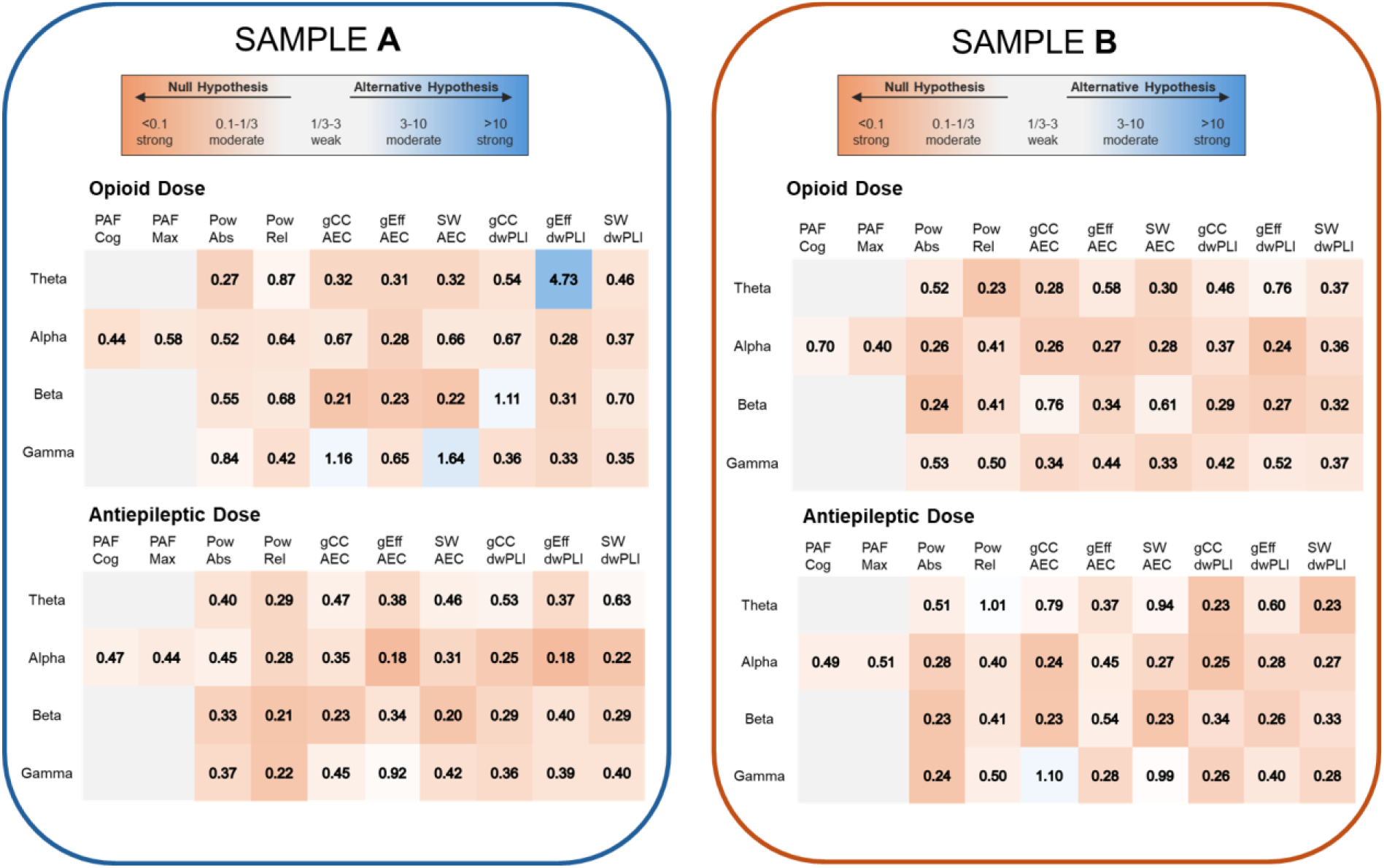
Dose-dependent effects of centrally acting analgesics on EEG parameters. Overview of BF10 resulting from regression models investigating dose-dependent effects of opioids (daily morphine equivalents; upper matrices) and antiepileptics (daily pregabalin equivalents, bottom matrices). For all models, age, pain intensity, and depressive symptoms were included as covariates. BF=Bayes Factor, PAFCog=peak alpha frequency (Center of Gravity), PAFMax=peak alpha frequency (Local Maximum), PowAbs=Absolute Power, PowRel=Relative Power, gCC=global Clustering Coefficent, gEff=global Efficiency, SW=Smallworldness, AEC=Amplitude Envelope Correlation, dwPLI=debiased weighted Phase Lag Index.

Thus, we did not find consistent evidence for, but mostly against, dose-dependent effects of centrally acting analgesics on rsEEG biomarker candidates of CP.

#### 3.2.3. Cumulative effects of analgesics

Lastly, we assessed the effects of cumulative analgesic use (including peripheral agents) measured by the MQS. In both samples, we found evidence against effects in most models (see Fig. 5). However, in sample A, we found strong evidence for some relationships between AEC-based network measures in the theta band and MQS scores. We specifically observed negative relationships between global clustering coefficient (*BF*_*10*_ *= 11*.*58*) and smallworldness (*BF*_*10*_ *= 9*.*73*) on the one hand and MQS scores on the other hand. Furthermore, we found strong evidence for a positive relationship between global efficiency (*BF*_*10*_ *= 10*.*73*) and MQS scores. Again, these findings were not replicated in sample B.

**Fig. 5:**
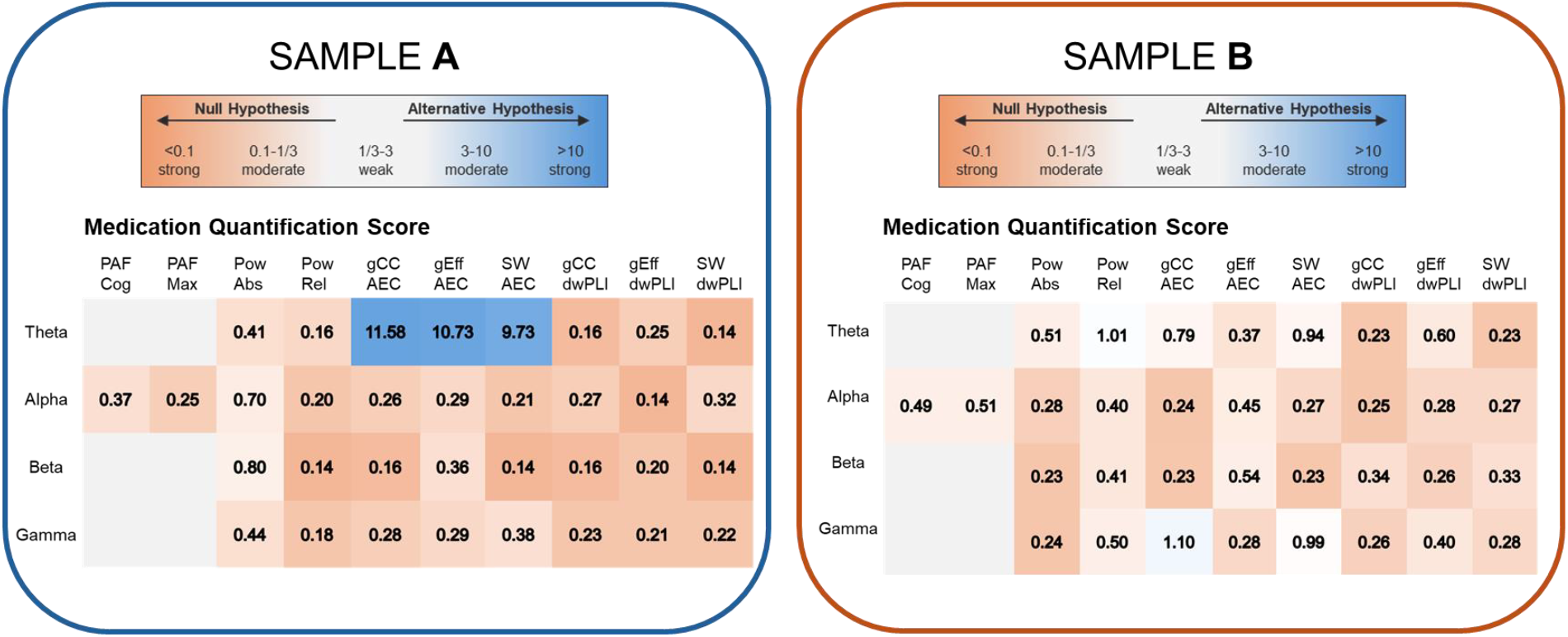
Cumulative effects of analgesics on EEG features. Overview of BF10 resulting from regression models investigating effects of scores on the MQS-4 on rsEEG biomarker candidates. For all models, age, pain intensity, and depressive symptoms were included as covariates. BF=Bayes Factor, PAFCog=peak alpha frequency (Center of Gravity), PAFMax=peak alpha frequency (Local Maximum), PowAbs=Absolute Power, PowRel=Relative Power, gCC=global Clustering Coefficent, gEff=global Efficiency, SW=Smallworldness, AEC=Amplitude Envelope Correlation, dwPLI=debiased weighted Phase Lag Index.

Thus, we mostly found evidence against cumulative effects of centrally acting analgesics on rsEEG biomarker candidates of CP.

## 4. Discussion

In this preregistered study, we analyzed the effects of centrally acting analgesics on rsEEG biomarker candidates of CP. To this end, we conducted Bayesian analyses of the effects of different analgesics on rsEEG parameters controlling for age, pain intensity, and depressive symptoms in two independent, large samples of people with CP (combined *n =220*). We almost exclusively found evidence against OPD, AED, and AD effects on rsEEG biomarker candidates of CP. These findings contribute to the development of rsEEG biomarkers in CP by helping to interpret previous and future studies in the field.

Previous studies on the EEG effects of analgesics in healthy people have shown that OPD increases delta and theta power [19]. In contrast, studies in people with CP have shown increases in alpha and beta power [9]. Research on the effects of AEDs in both healthy individuals as well as people with CP has demonstrated increased theta power and slowed peak alpha frequency [1; 19]. However, our analyses in two large samples of people with CP did not confirm these findings. While we did observe scant effects in the theta band, these were not replicable across samples. However, this, in principle, aligns with the aforementioned findings in healthy participants and supports the notion that changes in brain activity at theta frequencies play a role in the pathophysiology of CP [35].

Our study has several unique features. First, we performed Bayesian analyses, which enables quantification not only of evidence in favor of but also against effects of analgesics on EEG activity. Second, we controlled for important confounds of EEG activity by incorporating age, pain intensity, and depression scores as covariates. Third, we investigated the replicability of findings by performing all analyses in two independent large samples of people with CP recorded with different EEG methodologies. While our results cannot rule out any effect of centrally acting analgesics on rsEEG, they do indicate that centrally acting analgesics do not have a major impact on rsEEG measures in people with CP.

Several limitations apply to the present study. First, even though our samples are comparatively large for EEG studies in clinical populations, it might require larger sample sizes to detect subtle effects. A power analysis indicates that with a sample size of *n=118* for an ANCOVA with 2 groups and 3 covariates, we were able to detect an effect size of *f=0*.*26* (medium effect) with 80% power at a significance level of *alpha=0*.*05*. Thus, our study was not sufficiently powered to assess smaller than medium effects. Second, mostly Caucasian men and women were included, and all study participants were recruited from a single study center. Thus, the overall sample is not necessarily representative of other cohorts of people with CP. Third, although the clinical heterogeneity of our samples might reflect clinical reality, analyses in more homogeneous samples (e.g., only people with neuropathic pain) might yield different findings. Finally, the suitability of dry electrodes used in Sample B for analyzing higher frequency ranges like the gamma band is uncertain [15]. This limits the interpretability of gamma band findings in Sample B.

To draw conclusions for future studies, we mainly found evidence against effects of analgesics on common EEG features in people with CP. However, we still recommend reporting and considering analgesic use, especially when investigating lower frequency bands and participants using multiple substances. One option is to use the MQS as an easy-to-use tool to control for possible influences of analgesics on EEG activity. Furthermore, larger studies are needed to draw more definite conclusions about specific substance classes. Additionally, to establish rsEEG features as a reliable and valid biomarker of CP, it is crucial to consider other potential confounds of EEG activity in people with CP in future studies. Amongst others, these might include comorbid psychopathology and demographics.

In summary, our study provides evidence against effects of commonly used analgesics on peak alpha frequency, oscillatory power, and connectivity-based network metrics in two independent large samples of people with CP. This finding strengthens the validity of EEG biomarker candidates of CP and facilitates the interpretation of previous and future studies.

## Acknowledgments

The study and analysis plan were preregistered under https://osf.io/uzgxw. The authors thank the participants of the study and the members of the PainLabMunich for helpful discussions and comments on the manuscript. The study was supported by the TUM Innovation Network Neurotechnology in Mental Health (NEUROTECH) and the TUM School of Medicine Clinician Scientist Program (KKF). The authors report no conflicts of interest.

